# PhosX: data-driven kinase activity inference from phosphoproteomics experiments

**DOI:** 10.1101/2024.03.22.586304

**Authors:** Alessandro Lussana, Evangelia Petsalaki

## Abstract

**Motivation:** The inference of kinase activity from phosphoproteomics data can point to causal mechanisms driving signalling processes and potential drug targets. Identifying the kinases whose change in activity explains the observed phosphorylation profiles, however, remains challenging, and constrained by the manually curated knowledge of kinase-substrate associations. Recently, experimentally determined substrate sequence specificities of human kinases have become available, but robust methods to exploit this new data for kinase activity inference are still missing.

**Results:** We present PhosX, a method to estimate differential kinase activity from phosphoproteomics data that combines stateof-the art statistics in enrichment analysis with kinases’ substrate sequence specificity information. Using a large phosphoproteomics dataset with known differentially regulated kinases we show that our method identifies upregulated and downregulated kinases by only relying on the input phosphopeptides’ sequences and intensity changes. We find that PhosX outperforms the currently available approach to perform the same task, and we therefore recommend its use for data-driven kinase activity inference.

**Availability and implementation:** PhosX is implemented in Python, open-source under the Apache-2.0 licence, and distributed on the Python Package Index. The code is available on GitHub (https://github.com/alussana/phosx).

## Introduction

Kinases are major drivers of intracellular signalling, their activity contributing to virtually all biological processes. As they are often deregulated in disease, they are prevalent targets of approved drugs (1). Current phosphoproteomics techniques enable the quantification of thousands of phosphosites, allowing for unbiased assessments of intracellular signalling states. However, only a small fraction of discovered human phosphosites (2) have at least one known upstream kinase (3), posing a challenge to the estimation of kinase activity from phosphoproteomics data. At the same time, 150 human kinases have no known substrate, despite all of them likely being essential for normal human function (4). Recently, an atlas of substrate sequence specificity for 303 human Serine/Threonine kinases has been built for the first time, by employing a cell-free method known as positional scanning peptide assay (PSPA) (5). This resource can be used to link kinases to their potential target phosphosites in a data-driven manner opening the door to the accurate inference of all Ser/Thr kinase activities, regardless of the prior knowledge associated with them, and considering full phosphoproteomics datasets as opposed to the limited number of phosphosites with upstream kinase annotations. It also allows phospho-priming to be systematically taken into account when evaluating the affinity of a kinase for a given substrate. As we envision substrate specificity data being soon available also for the human Tyrosine kinases, we anticipate data-driven differential kinase activity inference to become a powerful tool in cell signalling research and drug discovery, but robust and benchmarked computational methods to perform this task are still missing. To address this need we developed PhosX, a software package to perform differential kinase activity inference from phosphoproteomics experiments using substrate sequence specificity data and state-of-the art statistics in enrichment analysis. We benchmarked its performance in recovering expected changes in kinase activity, and we show that it outperforms the currently proposed method to perform the same task while requiring fewer arbitrary parameters.

## Methods

### A. Description of the PhosX method

#### A.1 Input format of phosphosites

PhosX takes in input a ranked list of phosphopeptides, according to the change in intensity detected in a mass spectrometry-based phosphoproteomics experiment (typically a log fold change). The peptides’ sequences should be 10 amino acids long, with the modified residue in the 6^th^ position. Undefined amino acids are represented by the character `‘_’`. Every other residue is represented by the corresponding 1-letter symbol according to the IUPAC nomenclature for amino acids and additional phosphorylated Serine, Threonine or Tyrosine residues are represented with the symbols `‘s’`, `‘t’`, or `‘y’`, respectively.

#### A.2 Position Specific Scoring Matrices

PhosX estimates the affinity between human kinases and phosphosites based on the substrate sequence specificity measured in a previous study (5) and encoded in Position Specific Scoring Matrices (PSSMs). A kinase PSSM is a 10 × 23 matrix containing amino acid affinity scores at each one of the 10 positions of the target phosphosite. The 6^th^ position corresponds to the modified residue and should have non-0 values only for Serine, Threonine (for Ser/Thr kinases), or Tyrosine (for Tyr kinases) residues.

#### A.3 Phosphosite scoring

For each kinase PSSM, a score is assigned to each phosphosite sequence *S* that quantifies its similarity to the PSSM. First, a “raw PSSM score” is computed as:

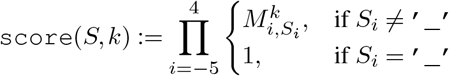

where *S*_*i*_ is the amino acid residue at position *i* of the phosphosite sequence *S*; 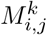 is the value of the PSSM for kinase *k* at position *i* for residue *j*. Raw PSSM scores for each kinase are then transformed between 0 and 1 based on the quantile they fall in, considering a background distribution of proteome-wide raw PSSM scores. These background scores are precomputed for each kinase on a set of more than 200, 000 human phosphosites annotated in the PhosphositePlus database (2) (September 2023). For each kinase, phosphosites with raw PSSM score equal to 0 are discarded, and the remaining are used to determine the values of the 10, 000-quantiles of the raw PSSM score distribution. The background 10, 000-quantiles raw PSSM scores for each kinase PSSM are used to derive the final PSSM scores for each phosphopeptide.

#### A.4 Running sum statistic

PhosX uses the PSSM scores to link kinases to their potential substrates. Each phosphosite is assigned as potential target to its *n* top-scoring kinases. In our benchmark we set *n* to its default value of 5. The activity change of a given kinase is estimated by calculating a running sum statistic over the ranked list of phosphosites, and by estimating its significance based on an empirical distribution generated by random permutations of the ranks. Let *C* be the set of indexes corresponding to the ranked phosphosites associated with kinase *k*; *N* the total number of phosphosites; *N*_*h*_ the size of *C*; *r*_*i*_ the value of the ranking metric of the phosphosite at rank *i*, where *r*_0_ is the highest value. Then, the running sum (*RS*) up to the phosphosite rank *n* is given by

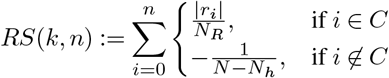

Where

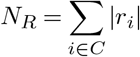

The kinase enrichment score (*ES*) corresponds to the maximum deviation from 0 of *RS* (6).

#### A.5 Permutation tests

For each kinase, PhosX computes an empirical *p* value of the *ES* by generating a null distribution of the *ES* through random permutations of the phosphosite ranks. A False Discovery Rate (FDR) *q* value is also calculated by applying the Bonferroni method considering the number of kinases independently tested. The number of permutations is a tunable parameter but we recommend performing at least 10^4^ random permutations to be able to compute FDR values *<* 0.05.

#### A.6 Kinase activity score

The activity score for a given kinase is defined as:

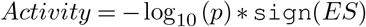

where `sign` is the sign function, and −*log*_10_(*p*) is capped at the smallest computable *p* value different from 0, *i.e*. the inverse of the number of random permutations. Activity scores greater than 0 denote kinase activation, while the opposite corresponds to kinase inhibition.

### B. Kinase activity inference with different methods

For each phosphoproteomics experiment, we used PhosX, GSEApy (7), and Kinex (8) separately, to infer the changes in kinase activities. GSEApy computes the same statistic as PhosX, but relies on prior knowledge to link kinases to their known substrates instead of using PSSM scores. We extracted such literature-derived kinase-phosphosite annotations for the human proteome from the Phosphositeplus database (2) and in each experiment we limited the analysis to kinases for which at least 4 annotated substrates appeared among the detected phosphopeptides, to ensure reliability of the results. Conversely, Kinex assigns substrates to kinases according to a PSSM score-based logic similarly to PhosX, but it imputes the changes in kinase activity based on a Fisher’s exact test assessing the enrichment of the substrates of a given kinase in the set of phosphopeptides having a log fold change more (for upregulation) or less (for downregulation) extreme than arbitrary thresholds. We set such thresholds at 0.5 and −0.5 to identify the log fold changes of phosphosites showing increased and decreased phosphorylation, respectively. As a metric for kinase activity we used the “Kinase activity score” in PhosX, the “Normalised enrichment score” (NES) in GSEApy, and the “Activity Score” in Kinex. In Kinex, if the reported “dominant direction” of the enrichment was “downregulated set”, we multiplied the Activity score by −1. To compare the estimated kinase activity changes across methods and experiments, for each method and for each experiment we scaled the inferred kinase activity values between 0 and 1 by first subtracting the minimum and then dividing by the maximum.

## Results

We developed PhosX, a software package to infer differential kinase activities from phosphoproteomics data that doesn’t require any prior knowledge database of kinase-phosphosite associations. PhosX assigns the phosphopeptides detected in an experiment to potential upstream kinases based on their substrate sequence specificity, which has been recently determined experimentally in the form of Position-Specific Scoring Matrices (PSSMs) (5); it then tests the enrichment of a kinase’s potential substrates in the extremes of the distribution of phosphopeptide intensity changes, using a KolmogorovSmirnov-like statistic analogous to the one implemented in the popular Gene Set Enrichment Analysis (GSEA) method (6) (Fig. 1A). PhosX can therefore be used to link observed changes in phosphorylation profiles to potential causal kinases, taking into consideration all detected phosphosites regardless of the amount of prior knowledge, if any, available for any specific phosphosite. PhosX is implemented in Python, can be easily installed from the Python Package Index, and can optionally take advantage of multiple cores to parallelise the computation.

**Fig. 1.**
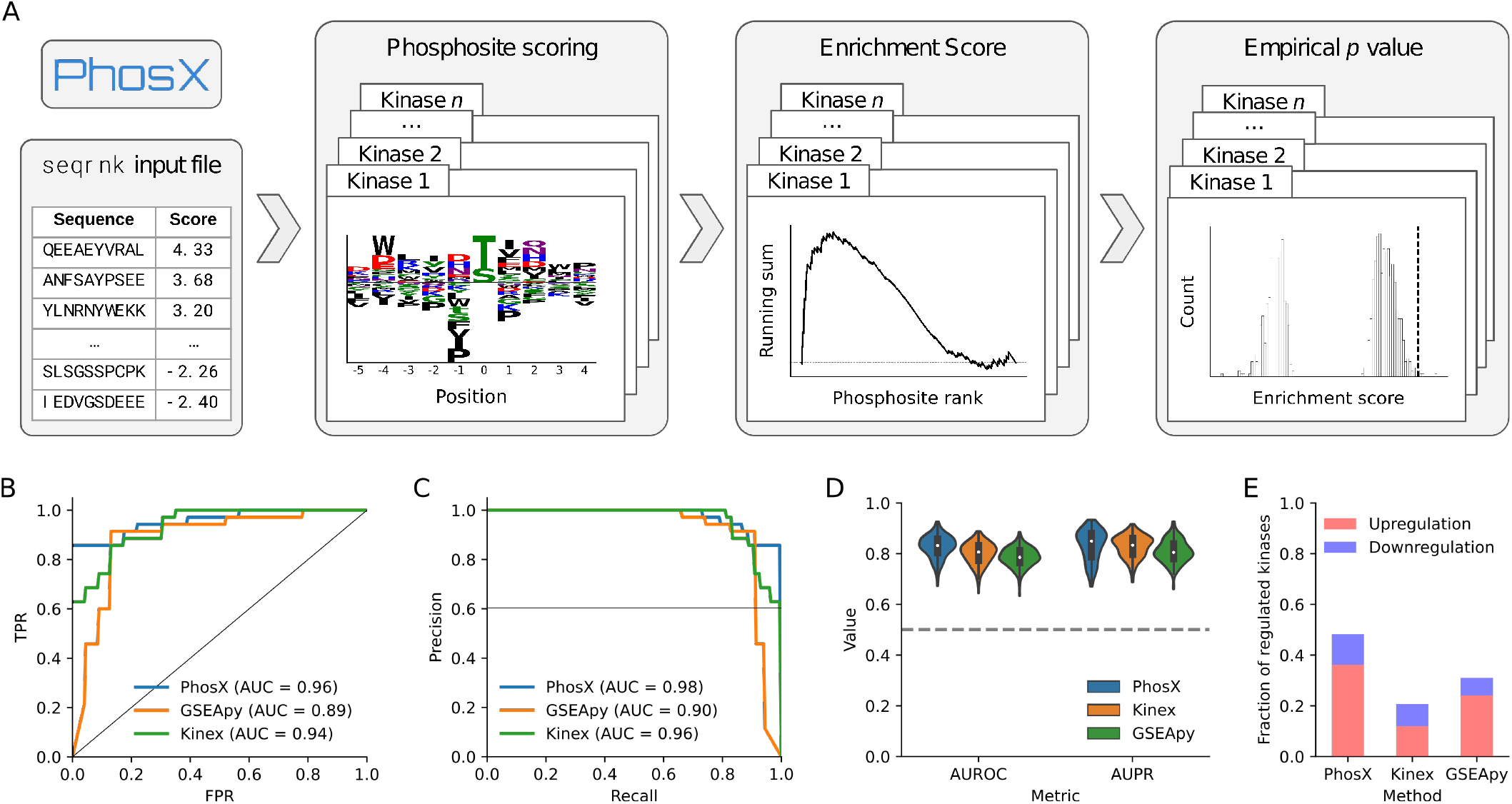
A. Schematic workflow of PhosX. PhosX takes in input a ranked list of phosphopeptides. Position Specific Scoring Matrices are used to score the input phosphosites and assign them to the most likely upstream kinases. The enrichment of a kinase’s substrates in the top or bottom phosphosite ranks is then tested via random permutations of the ranks to generate null distributions of an enrichment score statistic. **B**., **C**. Performance of the assessed methods in linearly separate upregulated from downregulated kinases in the benchmark dataset, measured by Receiver Operating Characteristic curves (B.) and Precision-Recall curves (C.). **D**. Performance of the assessed methods in linearly separate regulated kinases from a random negative set of kinases of equal size in the benchmark dataset. **E**. Fraction of instances of regulated kinases in the benchmark dataset found in the top-5 (upregulation) or bottom-5 (downregulation) percentiles of activity scores.

To evaluate the biological relevance of the kinase activity scores computed with PhosX, we collected a benchmark dataset of 94 perturbation experiments for which phosphosite log fold changes are publicly available (9). Each experiment is associated with kinases that are expected to be upor downregulated due to the nature of the perturbation (9). In our assessment we included two other methods with complementary characteristics: Kinex (8), which to our knowledge is the only currently available method inferring potential kinase-phosphosite interactions based on PSSM scores, but, differently from PhosX, it uses the Fisher’s exact test to predict kinase activity changes; and GSEApy (7), which, used in conjunction with the PhosphositePlus database (2), implements the same statistic as PhosX, but only considers known kinase-phosphosite interactions curated from the literature.

Because PSSMs for Tyrosine kinases are not available, PhosX and Kinex could compute activity scores for Ser/Thr kinases only. At the same time, GSEApy predictions were limited by the overlap between existing phosphosites annotations and the phosphopeptides detected in the experiments, leading to a consistently lower number of kinases for which an activity change could be estimated. To obtain comparable performance metrics, we considered the intersection of kinase activities that could be computed by all three assessed methods in the benchmark dataset. First, the linear separation based on the kinase activity score between positively- and negatively-regulated kinases in the ground-truth set was marginally greater for PhosX both when considering the area under the Receiving Operating Characteristic curve (AUROC) (Fig. 1B) and the area under the Precision-Recall curve (AUPR) (Fig.1C), indicating the ability to tell apart upfrom downregulation in the set of known regulated kinases. We then considered the task of distinguishing known regulated kinases from an equally sized set of examples where the kinases are not expected to be affected by the perturbations. We built such a negative set by randomly drawing kinase-experiment pairs, and repeated the evaluation 100 times, each time with a different random negative set, to account for technical variability. While we expect the presence of false negatives in this classification task, we observed favourable AUROC and AUPR for PhosX compared to the other methods (Fig. 1D). Finally, we computed the fraction of upregulated and downregulated ground-truth examples found in the respective extremes of the kinase activity score distribution according to each method. This metric provides a lower bound on the expected probability of discovering an upregulated or downregulated kinase among the ones inferred to be most affected by the perturbations. We defined such extremes as the top-5 and bottom-5 percentiles of activity score, and found that PhosX recovered substantially more kinases among the ones expected to be strongly regulated than the other methods (Fig. 1E). Conversely, Kinex had the worst performance in this task, with only rare instances of known regulated kinases being found at the extremes of the activity score distribution (Fig. 1, Suppl. Fig. 1), and we observed this being the case independently from the choice of log fold change threshold (Suppl. Fig. 2).

## Discussion and future work

We present PhosX, a tool to infer kinase activities from phosphoproteomics datasets based on the data and PSSM-based kinase substrate specificities alone. Despite our benchmarking being based on highly studied kinases with several known substrates, PhosX still outperformed GSEApy highlighting the power of using the unbiased PSSMs for this task. Similarly, despite using the same PSSMs as a basis, PhosX outperformed Kinex, in particular in identifying the correct regulated kinases in the extreme 5th percentiles than both methods, confirming the power of the rank sum statistic for this task (9). The statistical framework implemented in PhosX may be used together with any scoring function that assigns kinases to target phosphosites. While we can conclude that PSSM scores are useful for this purpose, substrate sequence specificity is not the only driver of kinase-phosphosite interactions. Presence of adaptor proteins, subcellular localisation, post-translational modifications, kinase expression, etc, are all factors that determine whether the phosphorylation of a target by a given kinase is mechanistically possible, but are not taken into account by the PSSM model alone. Furthermore, PSSMs of evolutionarily related kinases tend to be similar, leading to correlations in estimated kinase activities which may not be true. These limitations might be greatly alleviated by introducing better scoring functions, for example using machine learning classifiers that consider features beyond substrate specificity. Phosphoproteomics readouts result from the combined effect of the activity of all kinases and phosphatases, while the approach presented here treats kinases independently. We hypothesise that statistical modelling approaches that allow us to infer the activity of every kinase in combination with all the others may be worth exploring in future research. Finally, because we expect PSSMs to be generated also for the human Tyrosine kinases, we plan to integrate them in PhosX in a future update, increasing the usefulness of our tool by unlocking the data-driven kinase activity inference for the whole human kinome.

## Availability and implementation

PhosX is implemented in Python, open-source under the Apache-2.0 licence, and distributed on the Python Package Index (PyPI). The code is available on GitHub (https://github.com/alussana/phosx) and on Zenodo (DOI: 10.5281/zenodo.10868313). The results presented in this paper are fully reproducible through the Nextflow workflow available at https://github.com/alussana/phosx-benchmark.

## ACKNOWLEDGEMENTS

This work was supported by the European Molecular Biology Laboratory (EP, AL), and the EMBL International PhD Programme (AL). EMBL IT Support is acknowledged for provision of computer and data storage servers. The authors would like to thank Sophia Müller-Dott and Julio Saez-Rodriguez for fruitful discussions and for helping with the collection of the phosphoproteomics data used in the benchmark.

---

**Suppl. Fig. 1.**
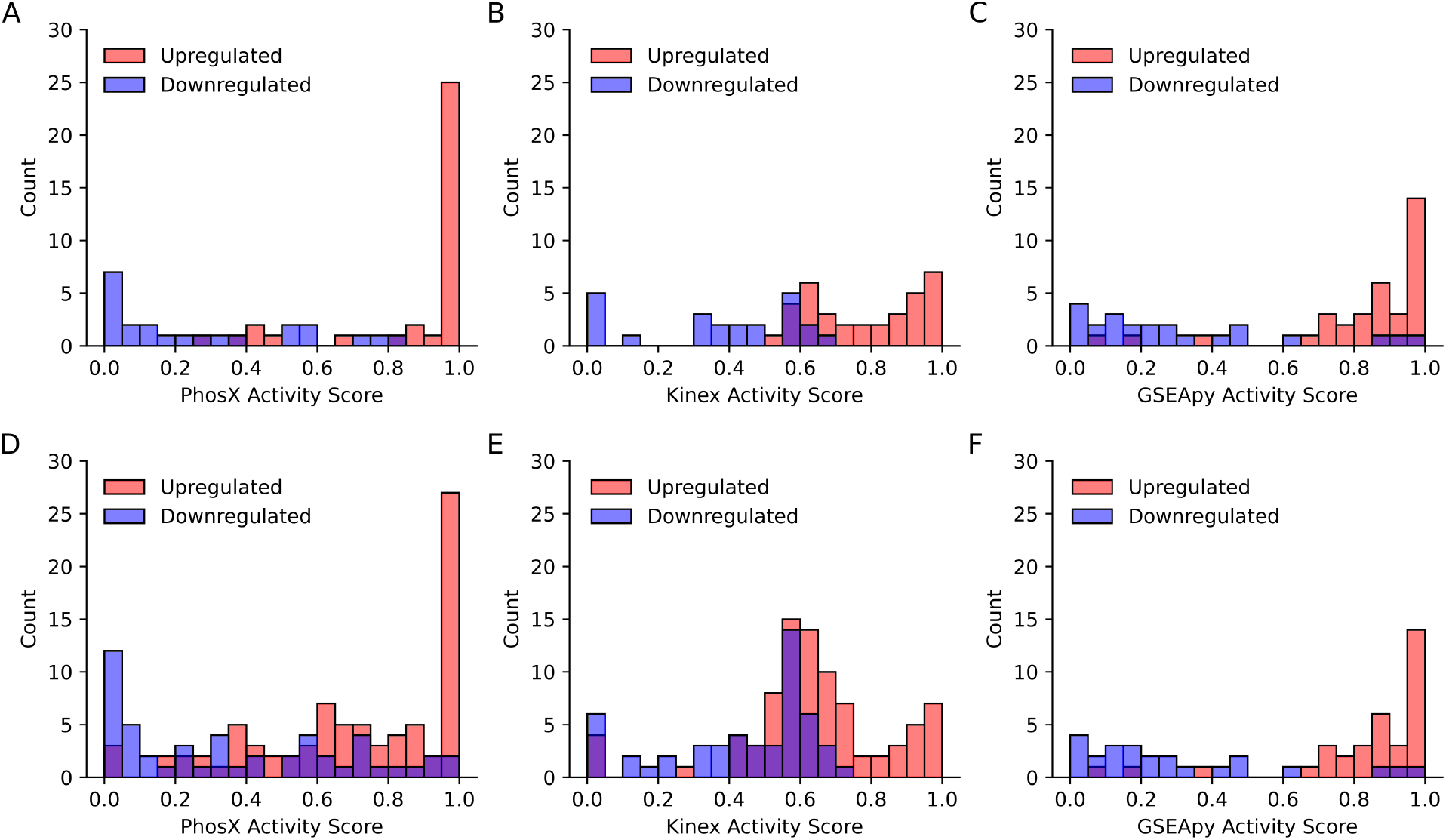
Distributions of the scaled kinase activity scores computed by PhosX (**A**., **D**.), Kinex (**B**., **E**.), and GSEApy (**C**., **F**.). Bins have a width of 5 percentiles. **A.-C**. Instances of upor downregulated kinases in the benchmark dataset for which a change in activity could be imputed by all the three methods are reported. **D.-F**. All instances of upor downregulated kinases in the benchmark dataset for which a change in activity could be imputed by the given method are reported.

**Suppl. Fig. 2.**
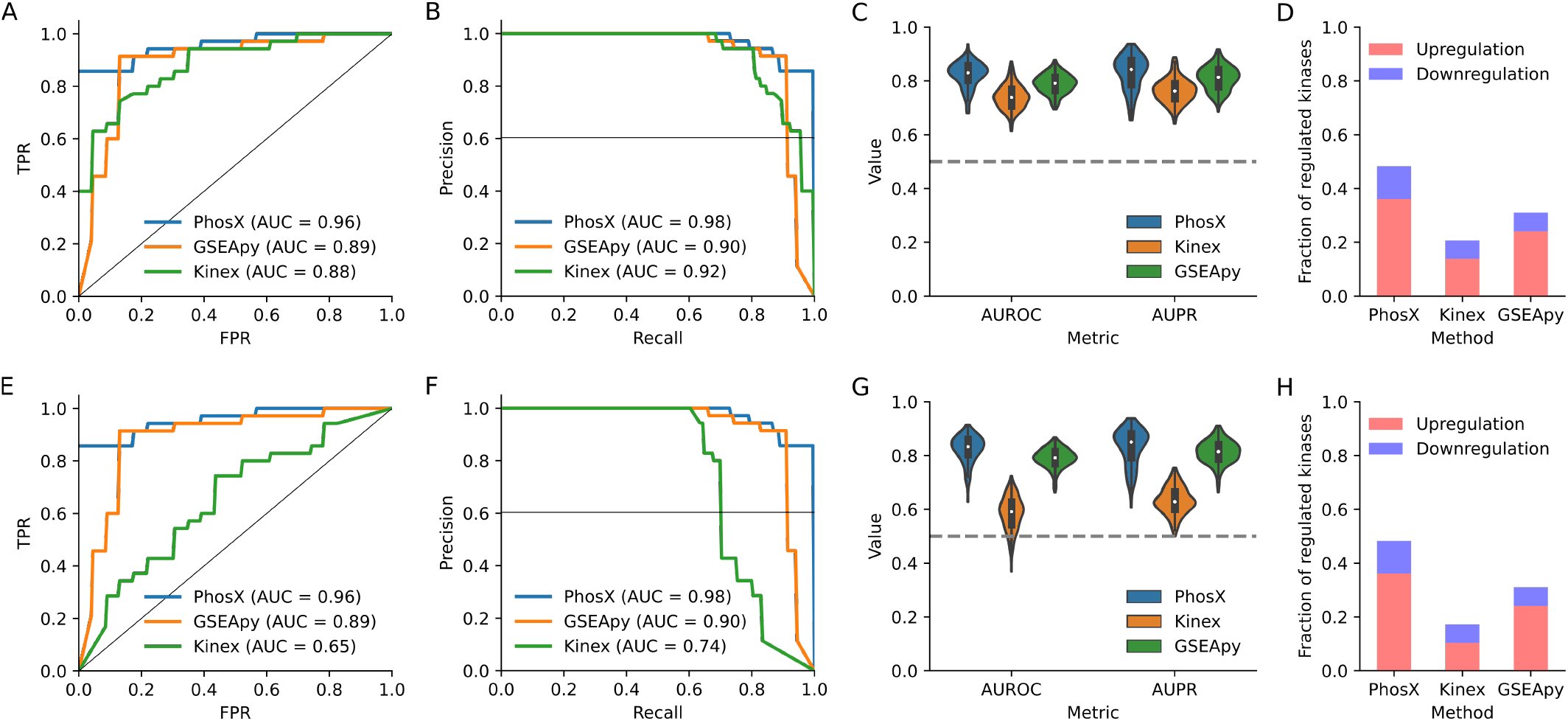
Benchmark results using alternative parameter values for Kinex’s log fold change (LFC) thresholds. **A.-D**. Figures refer to Kinex’s LFC = (1, *−*1). **E.-H**. Figures refer to Kinex’s LFC = (1.5, *−*1.5). **A**., **B**., **E**., **F**. Performance of the assessed methods in linearly separate upregulated from downregulated kinases in the benchmark dataset, measured by Receiver Operating Characteristic curves (A, E.) and Precision-Recall curves (B., F.). **C**., **G**. Performance of the assessed methods in linearly separate regulated kinases from a random negative set of kinases of equal size in the benchmark dataset. **D**., **H**. Fraction of instances of regulated kinases in the benchmark dataset found in the top-5 (upregulation) or bottom-5 (downregulation) percentiles of activity scores.

